# Paired stimulation for spike-timing dependent plasticity quantified with single neuron responses in primate motor cortex

**DOI:** 10.1101/2022.05.04.490684

**Authors:** Richy Yun, Jonathan H. Mishler, Steve I. Perlmutter, Eberhard E. Fetz

## Abstract

Spike-timing dependent plasticity (STDP) is an extensively studied topic. Previous studies have demonstrated stimulus induced targeted STDP both *in vitro* and *in vivo*, but a more consistent and robust method is required. We hypothesized there were two reasons contributing to the inconsistent results previously reported: 1. the measure of connectivity is poorly understood, and 2. the timing of stimulation is static or has low temporal specificity. To test our hypotheses, we applied paired stimulation to the primary motor cortex of awake primates. Single unit responses to stimulation were used as measures of connectivity, and we applied inter-stimulus intervals (ISIs) from ±0.1 to ±50 ms with sub-millisecond intervals. The excitatory single unit response resulted in very consistent changes after conditioning that was dependent on the ISI. Negative ISIs resulted in depression similar to classic STDP, but positive ISI also often resulted in depression. Normalizing the ISIs to the timing of the excitatory response revealed that potentiation only occurred if the second stimulus arrived before the response. Stimuli occurring around the time of the response often resulted in depression as strong as negative ISIs. We additionally tracked the changes in cortico-cortical evoked potentials (CCEPs), a commonly used measure of connectivity in plasticity experiments. CCEP changes showed a similar but more variable dependence to ISI. These results show that the classic STDP curve may be more difficult to induce due to interactions between excitatory and inhibitory circuitry, and that CCEPs may not be the ideal measure of changes in strength of connectivity.

## Introduction

Since the proposal of Hebbian plasticity, neuroplasticity has been an extensively studied topic (Hebb, 1949). Seminal *in vitro* studies expanded upon Hebb’s theory by showing that the specific timing of the pre- and postsynaptic cell activation is crucial for the changes in the synapse, called spike-timing dependent plasticity (STDP); if the postsynaptic cell is activated after the presynaptic cell the synapse is strengthened, and if the postsynaptic cell is activated before the presynaptic cell the synapse is weakened (Bi & Poo, 1998; Markram et al., 1997). Subsequent *in vivo* studies delivered activity-dependent stimulation, in which the postsynaptic site is stimulated immediately following spontaneous activity in the presynaptic site, to increase synaptic strength between cortical or cortico-spinal sites in both rodents and non-human primates (Jackson et al., 2006; McPherson et al., 2015; Nishimura et al., 2013; Rebesco et al., 2010).

Another method of investigating STDP is with paired stimulation, in which both the pre- and postsynaptic sites are directly activated via individual stimuli with a delay called the inter-stimulus interval (ISI). An advantage of paired stimulation is that it can be used to activate the postsynaptic site before the presynaptic site and artificially weaken synapses, a paradigm that is not possible activity-dependent stimulation. Paired stimulation can also be delivered open loop, and does not require any recording, providing potentially simpler clinical applications. Inducing STDP with paired electrical stimulation has also been demonstrated *in vivo* in both rodents and non-human primates; however, the induced changes were often inconsistent across tested channels (Rebesco & Miller, 2011; Seeman et al., 2017; Werk et al., 2006). Previous studies also used a limited number of ISIs and required up to 72 hours of continuous stimulation consisting of at least tens of thousands of paired stimuli to produce measurable, lasting changes.

We hypothesized that two factors may be limiting our understanding of applying paired stimulation *in vivo*. First, electrical stimulation in the context of paradigms for inducing plasticity is typically considered to be excitatory, but several studies have shown that stimulus response consists of a short-term excitatory response immediately followed by a long-term inhibitory response that is often strong enough to completely silence the cell for up to hundreds of milliseconds (Butovas & Schwarz, 2003; Logothetis et al., 2010; Yun et al., 2022). Electrical stimulation activates a bundle of fibers that project to the recorded site that excites the recorded neuron but also activates the surrounding inhibitory circuitry resulting in GABAergic inhibition (Butovas et al., 2006). As the dynamics of single unit responses change at sub-millisecond intervals, a more comprehensive set of ISIs with higher specificity may be necessary to fully explore paired stimulation *in vivo*.

Second, plasticity studies *in vivo* typically use macroscopic metrics as a measure of connectivity between two cortical sites, such as motor evoked potentials, directional tuning, coherence, or cortico-cortical evoked potentials (CCEPs) (Jackson et al., 2006; Seeman et al., 2017; Yazdan-Shahmorad et al., 2018). CCEPs are generated by delivering a stimulus to the presynaptic site and calculating the stimulus triggered averages of the local field potential response at the postsynaptic site. Studies usually quantify the response into a single value using the peak-to-peak amplitude or the root-mean square, with a larger value signifying a stronger connection.

However, CCEPs are complex and multiphasic, and the underlying sources of various phases and amplitudes of the response are still unclear (Boyer et al., 2018; Keller et al., 2014; Prime et al., 2020). Thus, a more direct measure of connectivity may clarify the changes induced by paired stimulation and their underlying mechanisms.

This study aims to better understand the application of paired electrical stimulation *in vivo* for the induction of STDP. We tested ISIs within a range of ±0.1 to ±50 ms at sub-millisecond intervals between pairs of channels up to 1.2 mm apart with the Utah array in primate motor cortex. We used the probability of a single-pulse stimulus in the presynaptic site evoking a spike in the postsynaptic site as a measure of connectivity strength. CCEPs were measured concurrently to compare the results between different connectivity metrics. In addition, we investigated changes in strength of the inhibitory response and evaluated all changes over a period of 10 minutes after conditioning to elucidate the underlying mechanisms.

## Materials and Methods

### Experimental design

#### Implants and surgical procedures

Two pigtail macaque monkeys (*Macaca nemestrina*), J and L, were unilaterally implanted with the Utah array (Blackrock Microsystems, Salt Lake City, UT) in the hand region of primary motor cortex (M1). Each array was 4×4 mm in size and consisted of 96 Iridium oxide coated electrodes with 1.5 mm length and 400 µm interelectrode distance. All procedures conformed to the National Institutes of Health *Guide for the Care and Use of Laboratory Animals* and were approved by the University of Washington Institutional Animal Care and Use Committee.

Implantation of the array was guided by stereotaxic coordinates. A 1.5 cm wide square craniotomy was performed over the hand region of the primary motor cortex to expose the dura. Three sides of the exposed dura were cut to expose the cortex; a Utah array was implanted and the dura was sutured over the array. Two reference wires were inserted below the dura and two were inserted between the dura and the skull. The bone flap from the craniotomy was returned, held in place by a titanium strap screwed onto the skull with 2.5 mm x 6 mm titanium skull screws. A second smaller titanium strap was fastened to the skull to secure the wire bundle. The connector pedestal for the array was attached to the skull with eight titanium skull screws and the incision closed around the pedestal base. Additional skull screws were placed around the base of the connector pedestal and a thin coat of dental acrylic (methyl methacrylate) was applied to the skull between the screws and the connector base for additional stability. Animals received postoperative courses of analgesics and antibiotics.

#### Experiment timeline

The two stimulated sites – “Pre” and “Post” sites – were determined by choosing a pair of channels in which stimulating the Pre site reliably evoked spikes in the Post site during a 1-minute preliminary recording. Each experiment consisted of 3 epochs: 1) pre-test – 10 minutes of test stimulation delivered to the Pre site, 2) conditioning – 10 minutes of paired stimulation, and 3) post-test – 10 minutes of test stimulation delivered to the Pre site (Figure 1). In addition, 5 minutes of baseline recording with no stimulation was collected both immediately before and after each experiment. As the stimulation could have long-lasting effects, conditioning between each channel pair was limited to once per day. Animals were trained to calmly sit in a primate chair throughout the duration of each experiment.

**Figure 1.**
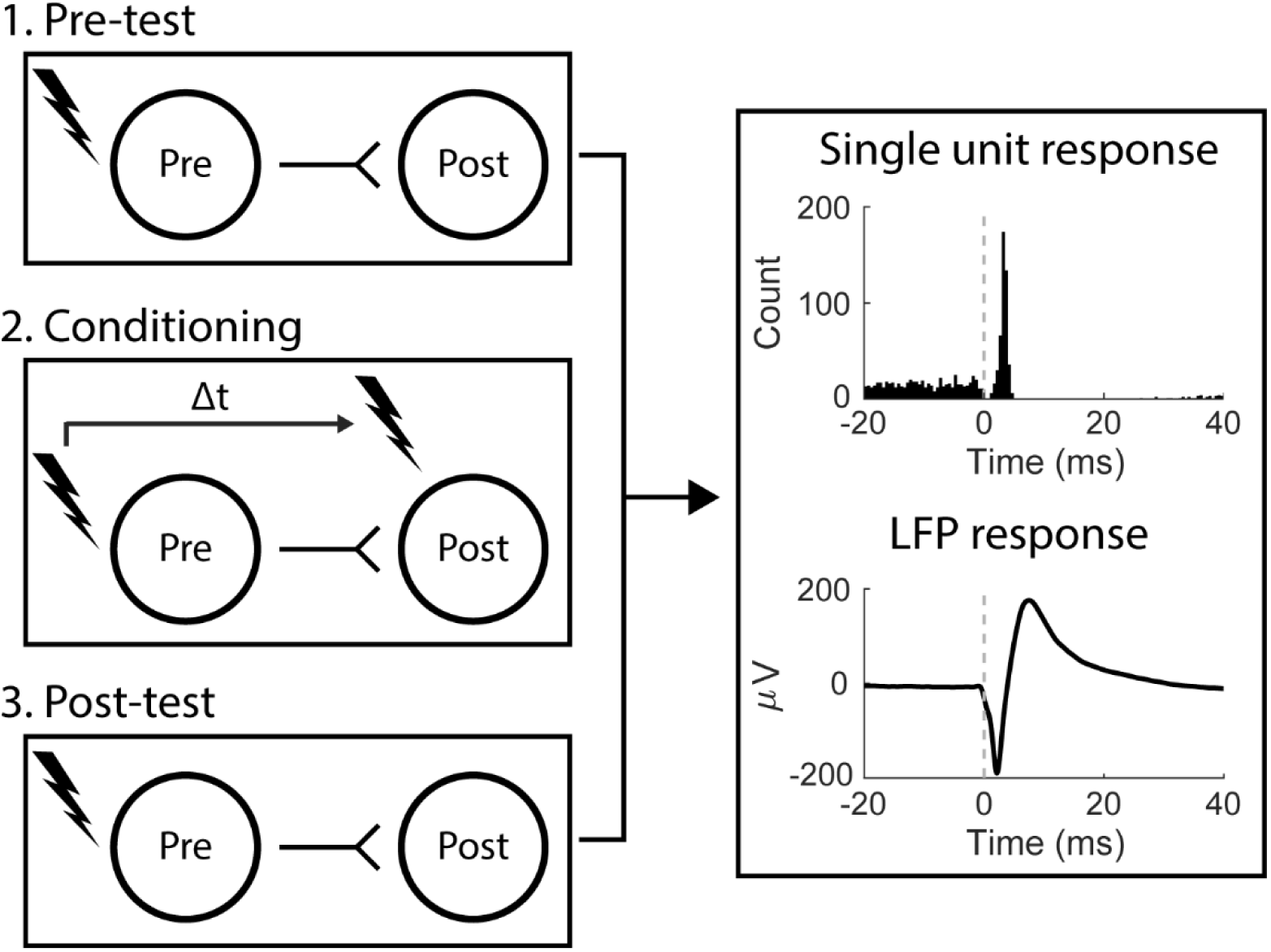
Experimental design. Each experiment consisted of three epochs – pre-test, conditioning, and post-test. During the test epochs stimuli were delivered to the presynaptic site and responses captured at the postsynaptic site. Conditioning consisted of paired stimulation between the pre- and postsynaptic sites at an arbitrary delay, Δt.

#### Cortical stimulation

All stimuli delivered were single-ended, single pulse with 200 μs phase width. Test stimuli were delivered at 5 Hz, Poisson distributed and paired stimulation during Conditioning were delivered at 10 Hz, Poisson distributed. These frequencies have been shown not to heavily affect the stimulus responses over time (Yun et al., 2022). The interstimulus interval (ISI) between the paired stimuli was randomly determined each day. All stimuli used an amplitude of 15 µA as previous studies showed it to be sufficient to reliably activate the stimulated site and generate responses in sites up to 4 mm away (Butovas & Schwarz, 2003; Hao et al., 2016; Yun et al., 2022). The stimulus intensity was kept the same across all sessions rather than adjusted to the stimulus response to limit current spread. We chose channel pairs in which stimulating one site evoked spikes in the other within 5 ms to ensure the presence of monosynaptic projections. The distance between channel pairs was between 400 and 1200 µm (up to 3 channels apart).

### Data analysis

#### Recording and spike sorting

We used the Neural Interface Processor (Ripple Neuro, Salt Lake City, UT) for both recording and stimulation. All 96 channels of the Utah array were recorded at 30 kHz throughout every experiment. Stimulus artifacts typically lasted 1-1.5 ms, and each stimulus pulse triggered a fast settle of 0.8 ms to minimize the stimulus artifact without affecting the stimulus evoked spike. The signals were bandpass filtered between 1 and 2 kHz for spike sorting to further minimize the artifact.

Spikes were sorted offline using custom MATLAB code through two-window discrimination; a window of -0.5 to 1.5 ms each time the signal crossed a negative threshold were collected, then two time-amplitude windows were manually selected to detect the peak and trough of the spike waveform. The investigator sorting the spikes were blind to the stimulus condition to ensure objective evaluation.

#### Stimulus evoked spikes and inhibitory response duration

Detailed methods of detecting stimulus evoked spikes and calculating the inhibitory response duration can be found in Yun et al. 2022. In brief, stimulus evoked spikes were found by calculating the peristimulus time histogram (PSTH) of each spike. To isolate the evoked spikes from the spontaneous activity, we defined upper and lower thresholds in the PSTH as the histogram mean plus or minus 2 times the standard deviation from -20 to -2 ms. We then found the largest peak in the PSTH from 1 to 15 ms after stimulation that was larger than the upper threshold and tracked adjacent bins in both directions until we reached the lower threshold on both sides. All spikes occurring within this window were denoted as stimulus-evoked spikes. If no peak was greater than the threshold the spike was not considered to have been evoked by stimulation. The average timing of evoked spikes was calculated as the median timing of evoked spikes from stimulation onset.

We measured the duration of inhibition by removing all evoked spikes and calculating the time between the onset of stimulation and the next spontaneous spike. Inhibition was deemed to be stronger when the delay from stimulation onset to the next spontaneous spike was longer. We discarded any stimuli for which the subsequent stimuli occurred before the next spontaneous spike.

#### Cortico-cortical evoked potential calculation

One method of calculating the magnitude of CCEPs is the peak-to-peak amplitude (Seeman et al., 2017; Zanos et al., 2018). However, as CCEPs can vary in timing and waveform the root mean square (RMS) for the area under the curve is sometimes used as a more consistent substitute measure of CCEP magnitude (Dionisio et al., 2019; Enatsu et al., 2013). However, as the changes were much more variable when using RMS or the area under the curve (Supplementary Figure 1), comparisons with single unit responses and the changes of measures over time used the peak-to-peak amplitude.

To obtain the peak-to-peak amplitude, stimulus triggered averages of raw LFPs were calculated from 10 ms before to 50 ms after stimulus onset. The amplitude was then calculated by subtracting the largest trough from the largest peak in a window of 1.5 to 50 ms after stimulation. To obtain the RMS magnitude we calculated the RMS over stimulus triggered averages of raw LFPs from 1.5 to 50 ms after stimulus onset:

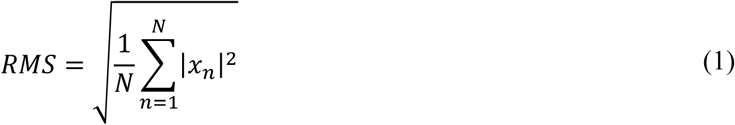

where *x* is the signal and *N* is the number of samples. The area under the curve was calculated as the sum of the rectified average response in a window of 1.5 to 50 ms after stimulation.

Stimulus artifacts were removed from all CCEP illustrations presented in the study by blanking out 0 to 1.5 ms from stimulation. If the calculated magnitude was less than 4 standard deviations of the signal from -10 ms to 0 ms of stimulus onset it was assumed there was no CCEP at the recorded site. CCEPs were not present in all channel pairs as they require raw local field potentials (LFPs) which are more susceptible to stimulus artifact.

#### Statistical analyses

Changes in stimulus response were calculated as the percent difference between the responses during the Pre- and Post-test epochs. Fisher’s exact test was used to compare the probability of evoking a spike before and after conditioning. Wilcoxon rank-sum test was used to compare changes in inhibition duration due to the nonparametric nature of the distributions. Student’s t-test was used to compare changes in CCEPs. Statistical tests used and p-values for significance are reported in individual analyses.

## Results

We delivered paired stimulation between pairs of sites in the motor cortex of two monkeys: 86 conditioning experiments and 35 control experiments: 37 conditioning experiments and 11 control experiments with 26 unique channel pairs in Monkey J, and 49 conditioning experiments and 24 control experiments with 14 unique channel pairs in Monkey L. We attempted to test the same channel pair with as many different experimental conditions as possible, but single units were often not stable for extended periods of time.

### Stimulus responses

Single pulse intracortical electrical stimulation reliably evoked spikes followed by a long inhibitory response (Figure 2A). We calculated the probability of evoked spikes by finding the peak of the excitatory response and the duration of inhibition by finding the timing of the spike following each stimulus that is not an evoked spike (*Stimulus evoked spikes and inhibitory response duration* in *Methods and Materials*). Channel pairs were chosen such that the evoked spike arrived within 5 ms to ensure the presence of monosynaptic projections, and the inhibitory response lasted between 10 and 100 ms (Figure 2B). As the stimulus intensity was kept consistently low across all sessions to limit current spread, there was a wide distribution of initial evoked probabilities (Figure 2B).

**Figure 2.**
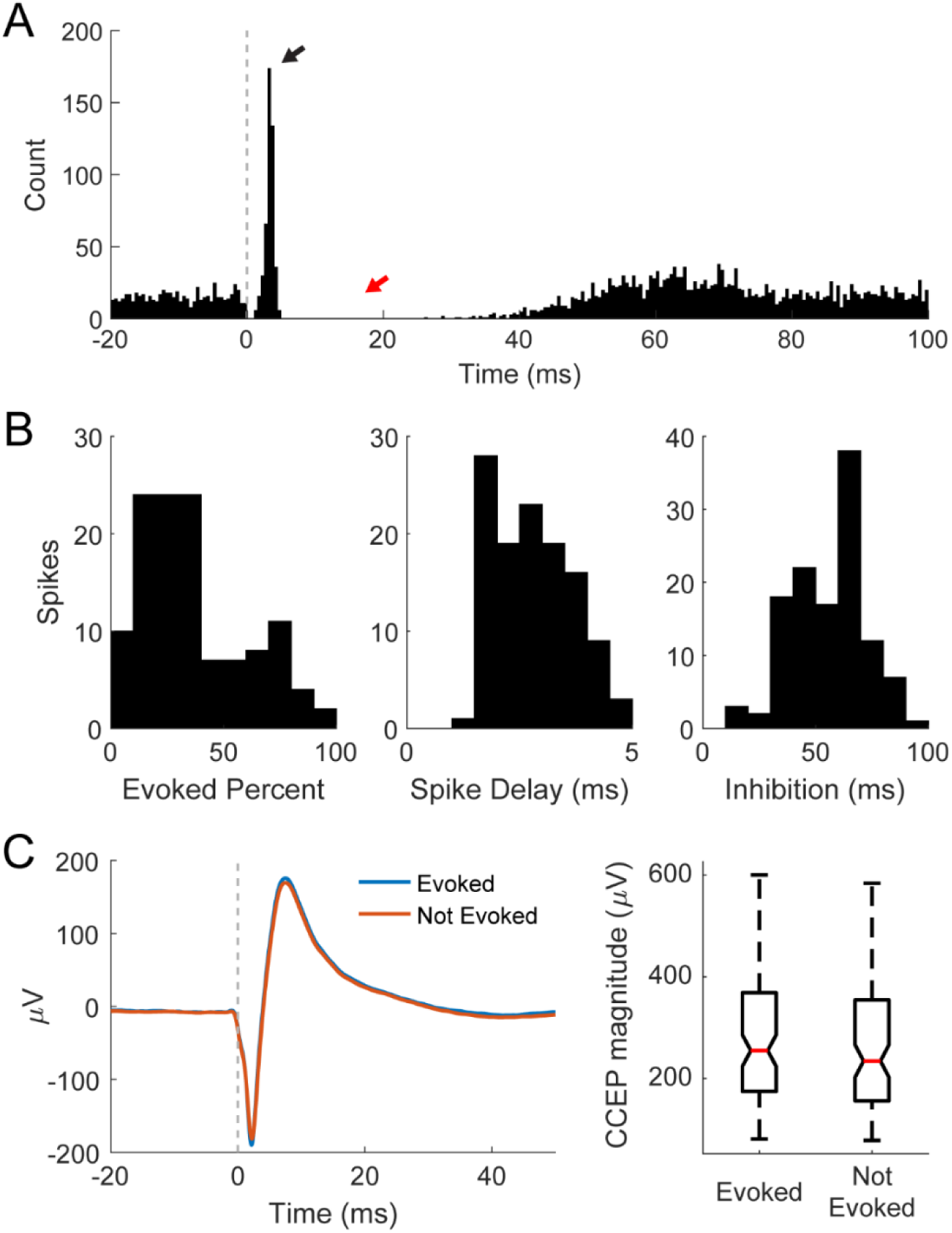
Stimulus responses. **A)** Example PSTH of a spike showing the stimulus evoked spikes occurring quickly after stimulus onset (black arrow) and the strong inhibitory response lasting 10s of milliseconds following the evoked spikes (red arrow). **B)** Distributions of preconditioning evoked spike percent, spike delay, and inhibition duration. **C)** Example CCEP trace. Example CCEPs showing similarity between CCEPs of stimuli that evoked spikes and those that didn’t (left). The same holds across all spikes throughout the experiment (right) showing the independence of the two measures.

We also simultaneously measured cortico-cortical evoked potentials (CCEPs) (Figure 2C). The responses typically had a narrow trough with the minimum value between 3 and 5 ms followed by a wider peak with the maximum value between 5 and 20 ms, similar to those seen in previous intracortical studies (Seeman et al., 2017; Zanos et al., 2018). The entire response typically lasted about 50 ms after stimulation. To ensure that evoked spikes were not contributing to the CCEPs, we calculated the CCEPs triggered from stimuli that evoked spikes or stimuli that did not evoke spikes (Figure 2C, left). There was no significant difference between the two CCEPs across all experiments, showing that they are independent measures (Figure 2C, right).

### Stimulus evoked spikes reflect STDP-like changes

Paired stimulation caused significant changes in the probability of evoking spikes before and after conditioning depending on the ISI (Figure 3A). Paired stimulation in which the Pre site was stimulated following the Post site (negative ISIs) resulted in a consistent decrease in the evoked spike probability. This depressive state typically occurred within a 20 ms time window and exponentially decayed with larger ISIs, similar to what is seen in classic spike-timing dependent plasticity (STDP).

**Figure 3.**
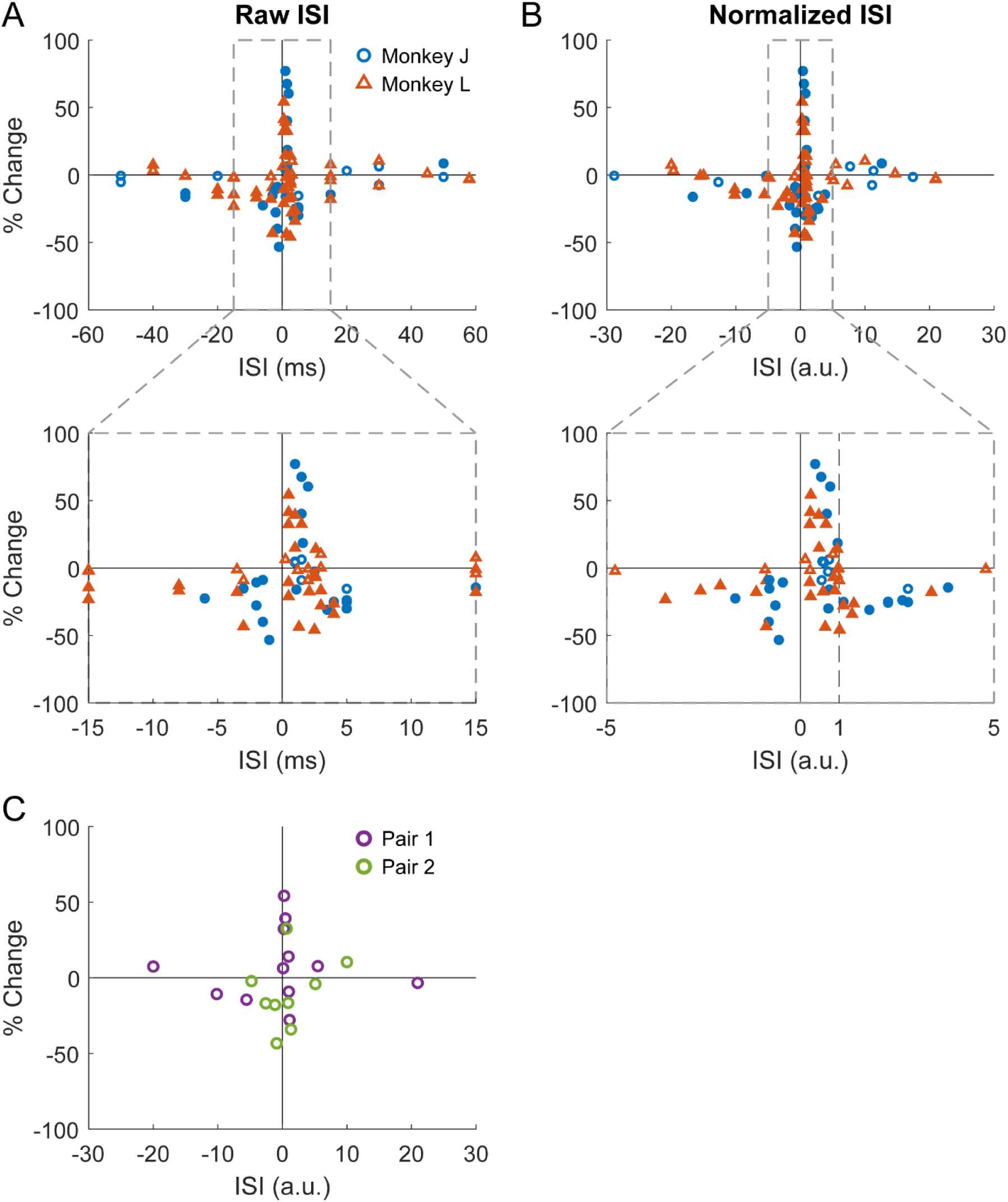
Evoked spike changes. Changes in evoked spike probabilities plotted against **A)** raw ISI and **B)** normalized ISI. Filled in data points are statistically significant from 0 (Fisher’s exact test, p<0.05). **C**. Changes in evoked spike probabilities of two pairs of presynaptic site and postsynaptic spike at different ISIs.

When the Pre site was stimulated before the Post site (positive ISI) with a very short ISI, there was typically potentiation, again similar to classic STDP. However, with ISIs > 1 ms there was high variability in the induced changes. All potentiation occurred with ISIs less than 3 ms, and ISIs of 3-20 ms typically resulted in depression almost as strong as negative ISIs, contradicting classic STDP. ISIs greater than +20 ms had little to no change.

To better understand the timing of the effects of paired stimulation, we additionally analyzed the changes by normalizing the ISI to the average timing of evoked spikes following test stimuli (Figure 3B). Positive ISI results became more consistent, with ISIs arriving before the evoked spike resulting in a greater change of an increase in evoked spike probability that reverses as it gets closer to the evoked spike timing. ISIs less than 0.5 of the evoked spike timing consistently led to potentiation, whereas ISIs immediately before and after the evoked spike timing led to depression. Plotting the same channel pairs tested at different ISIs show that the trend holds true; the different ISIs seem to drive the changes rather than the variability between channel pairs (Figure 3C). All data showing changes in responses is plotted against normalized ISI from this point forward.

We additionally analyzed whether there were consistent changes in baseline firing rate of the spikes before and after conditioning, as changes in excitability has been shown to affect both the probability of evoking a spike and the inhibitory response (Yun et al., 2022). However, there were no consistent changes in firing rate due to conditioning, allowing us to directly compare the measures between the pre- and post-test epochs (Figure 4).

**Figure 4.**
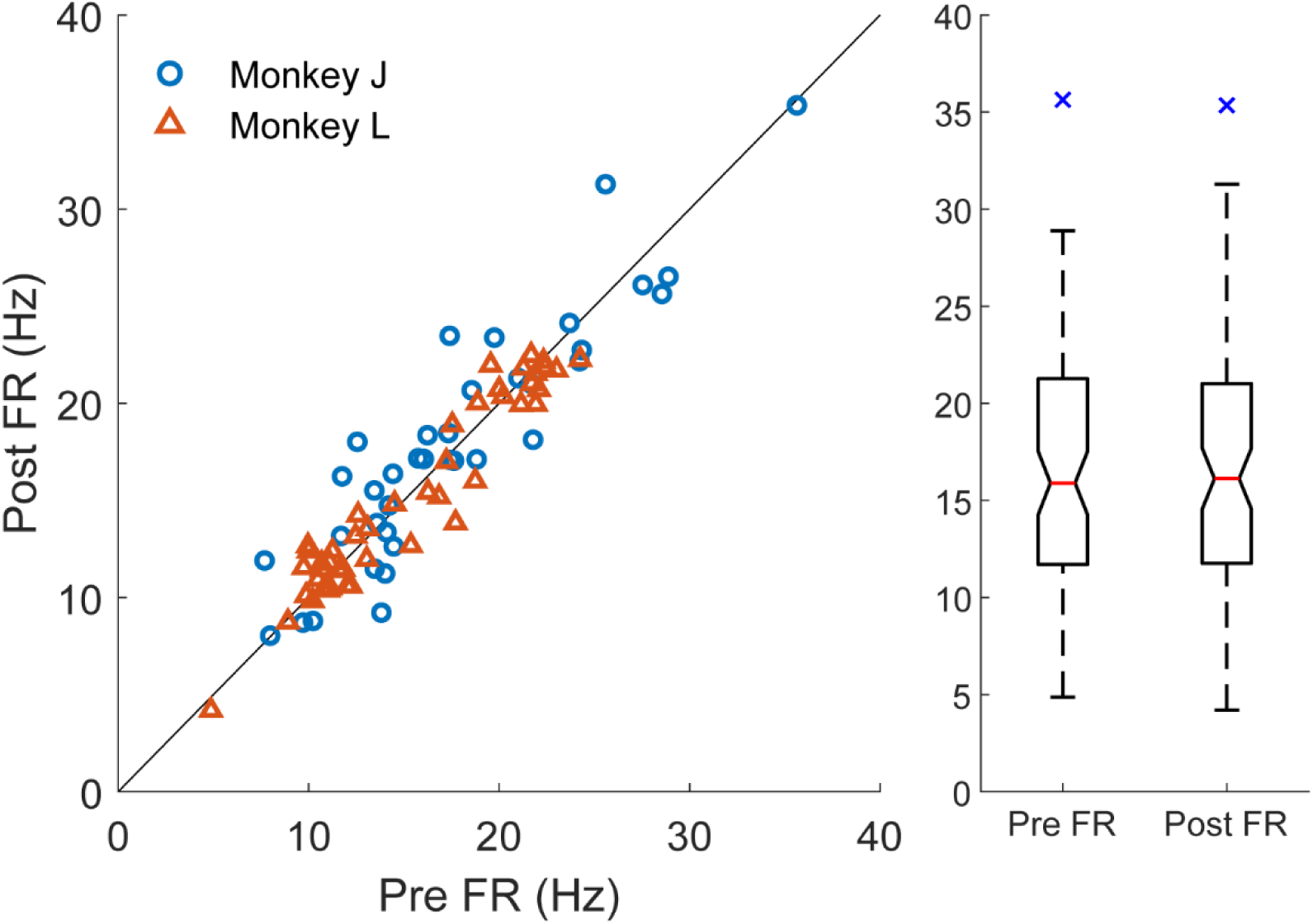
Firing rate. The firing rate of the spikes during the Pre- and Post-test epochs (left). The diagonal line shows y=x. There was no significant difference in firing rate between the epochs as shown in the boxplot on the right (p=0.79, Wilcoxon signed-rank test). The blue x shows outliers.

Finally, we also assessed whether the initial probabilities affected the highly variable changes observed at positive normalized ISIs between zero and 1 (second pulse arrives before the evoked spike timing). A strong initial response may be more difficult to increase due to saturation of activation, whereas a low initial response may be easier to increase. The initial probability of evoking a spike had a weak but significant negative correlation with the changes induced by conditioning, which may explain the high variability seen in that region of ISIs (Figure 5A).

**Figure 5.**
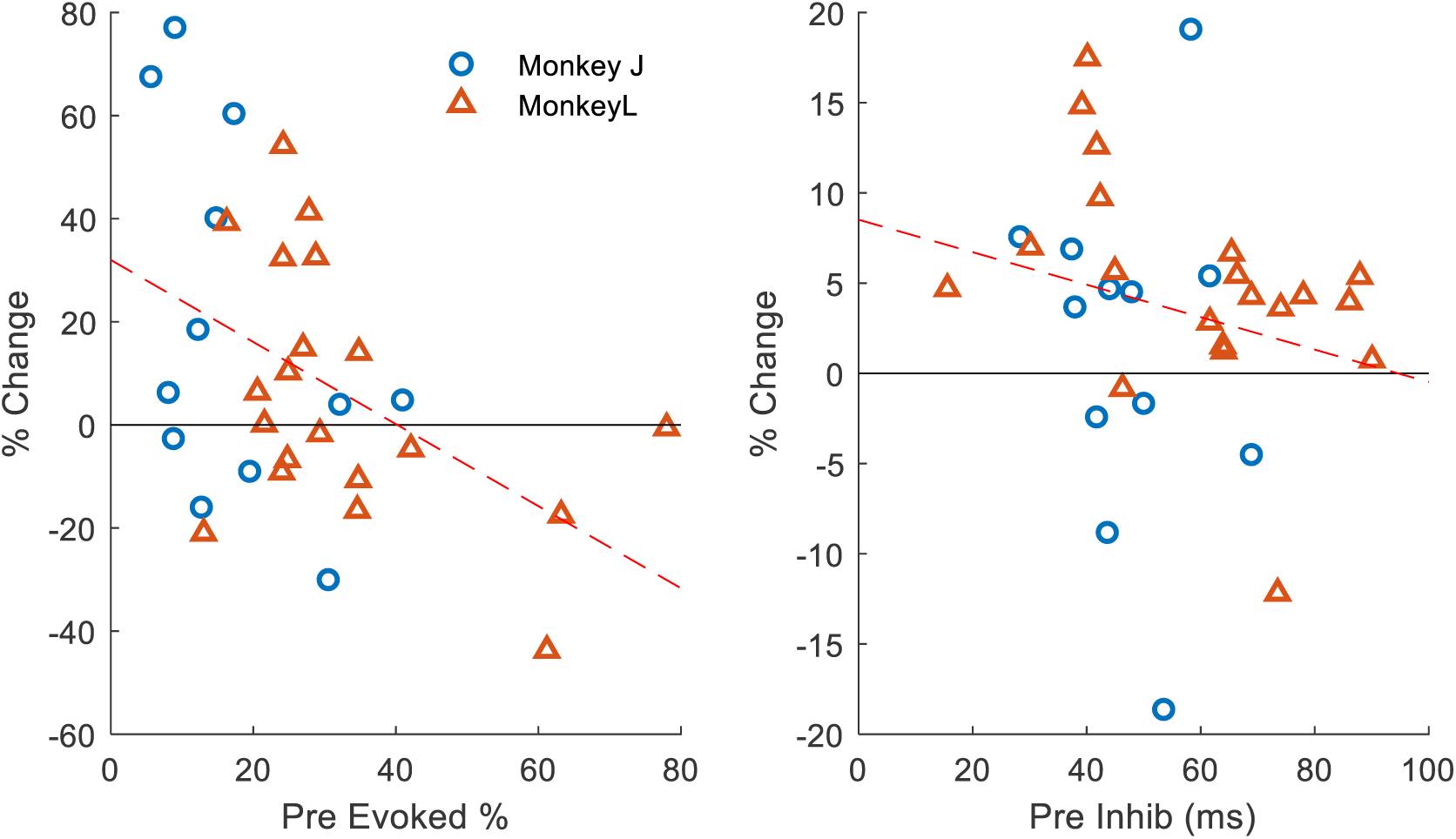
Evoked spike changes relative to initial evoked rate. **A**. Percent change in evoked spike probability with respect to raw evoked probability during the pre-test epoch for all sessions in which 0 < normalized ISI < 1. The dashed red line shows the linear regression (Pearson correlation, rho= -0.45, p=0.01). **B**. Percent change in inhibition duration with respect to inhibition duration during the pre-test epoch. The dashed line shows the linear regression (Pearson correlation, rho=-0.22, p=0.24).

### Inhibitory responses

We tracked the changes in the inhibitory response duration of single units following stimulation to elucidate whether the stimulation is causing changes in the inhibition circuitry. The term “inhibitory” as used here refers to the silencing of the neuron following the stimulus evoked excitatory response that is likely driven by local inhibitory circuitry. Figure 6 shows the percent change in the median inhibitory response after conditioning relative to normalized ISI. The changes were not as large or consistent as the changes in the probability of evoked spikes, and the high variability makes interpretation of the effects of ISI difficult. However, the magnitude of the changes loosely followed classic STDP, with smaller ISIs typically resulting in larger deviations, suggesting the conditioning was affecting the inhibitory response, and subsequently the local inhibitory circuitry.

**Figure 6.**
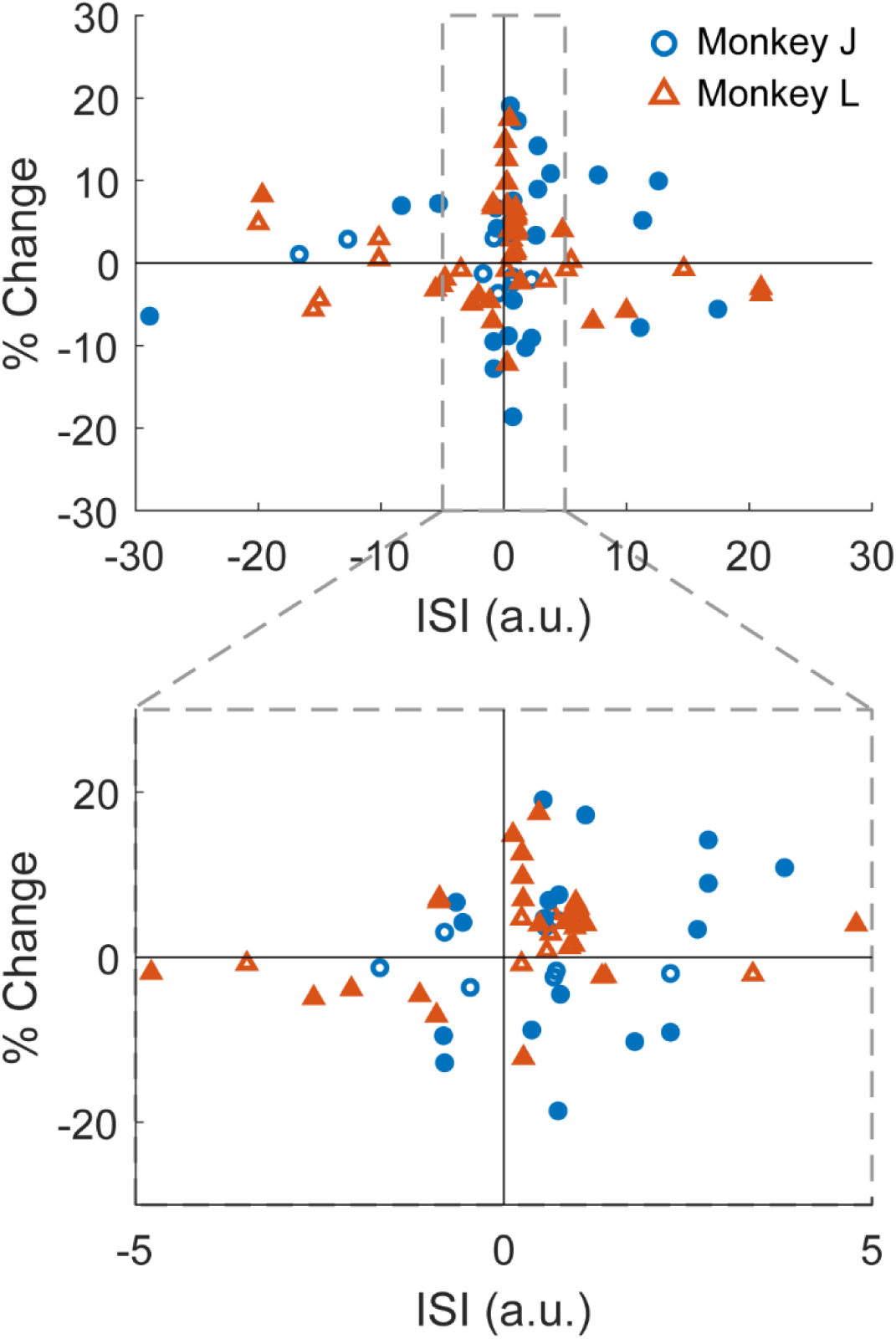
Inhibition changes. A. Raw ISI B. Normalized ISI. Filled in data points are statistically significant from 0 (Fisher’s exact test, p<0.05).

### CCEPs

We also tracked CCEPs before and after conditioning to compare the two measures of connectivity. Not every channel pair produced CCEPs as CCEPs are calculated with averaged LFP traces which are more susceptible to noise, especially with the proximity of channel pairs used in our experiments. The changes reflected in CCEPs were not as consistent as with evoked spikes relative to the ISIs of the paired stimulation (Figure 7). The negative stimulation condition and the positive stimulation condition that arrived before evoked spikes had similar trends as with evoked spikes but the positive stimulation that arrived after evoked spikes had much higher variation. A large number of changes across the entire ISI spectrum deviated from the classic STDP curve.

**Figure 7.**
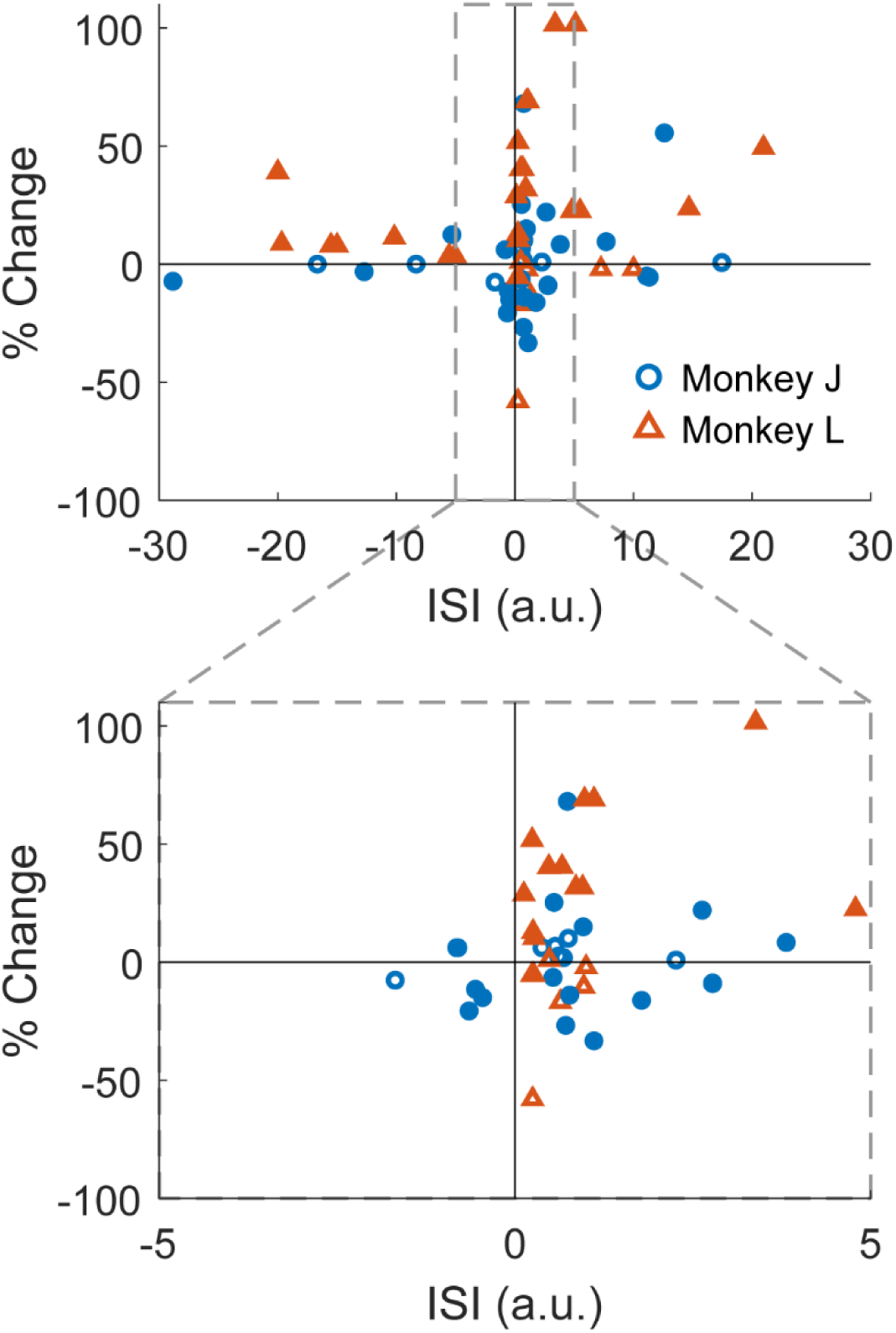
CCEP changes. CCEP changes relative to normalized ISI. Filled in data points are statistically significant from 0 (2 sample t-test, p<0.05).

We additionally plotted the changes in CCEPs as measured with root-mean squared or the area under the curve, as CCEPs are often complex and multiphasic (Supplementary Figure 1).

However, the changes in these two measures were extremely variable especially in experiments with small ISIs. In addition, peak-to-peak is the most commonly used metric of CCEPs when used for connectivity analysis. As a result, all following analyses are performed with CCEPs as measured with peak-to-peak.

### Comparisons between measures

We tested whether there was a direct relationship between the changes in evoked spike probability and inhibition for all ISIs but found no significant correlation (Figure 8). However, changes caused by different ISIs may be due to different mechanisms. As a result, we defined four distinct groups – 1) negative ISI greater than -20 ms, 2) positive ISI less than 10 ms with potentiation, 3) positive ISI less than 20 ms with depression, 4) all other ISIs (Figure 9A). We observed no significant correlation between the three measures in each of the four groups (Supplementary Figure 2).

**Figure 8.**
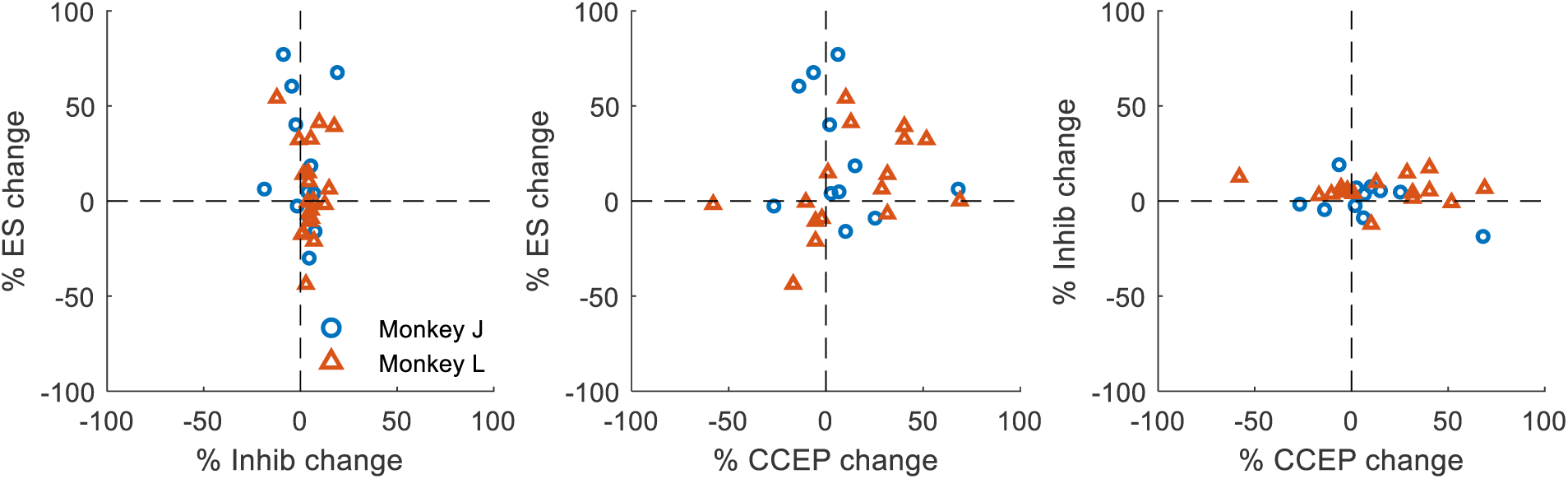
Correlation between evoked spikes, inhibition duration, and CCEP changes. There is no significant pairwise correlation between changes in evoked spike probabilities, inhibition duration, and CCEP amplitudes (Pearson correlation, rho= -0.13, p= 0.47; rho=0.11, p=0.57; rho=0.18, p=0.37).

**Figure 9.**
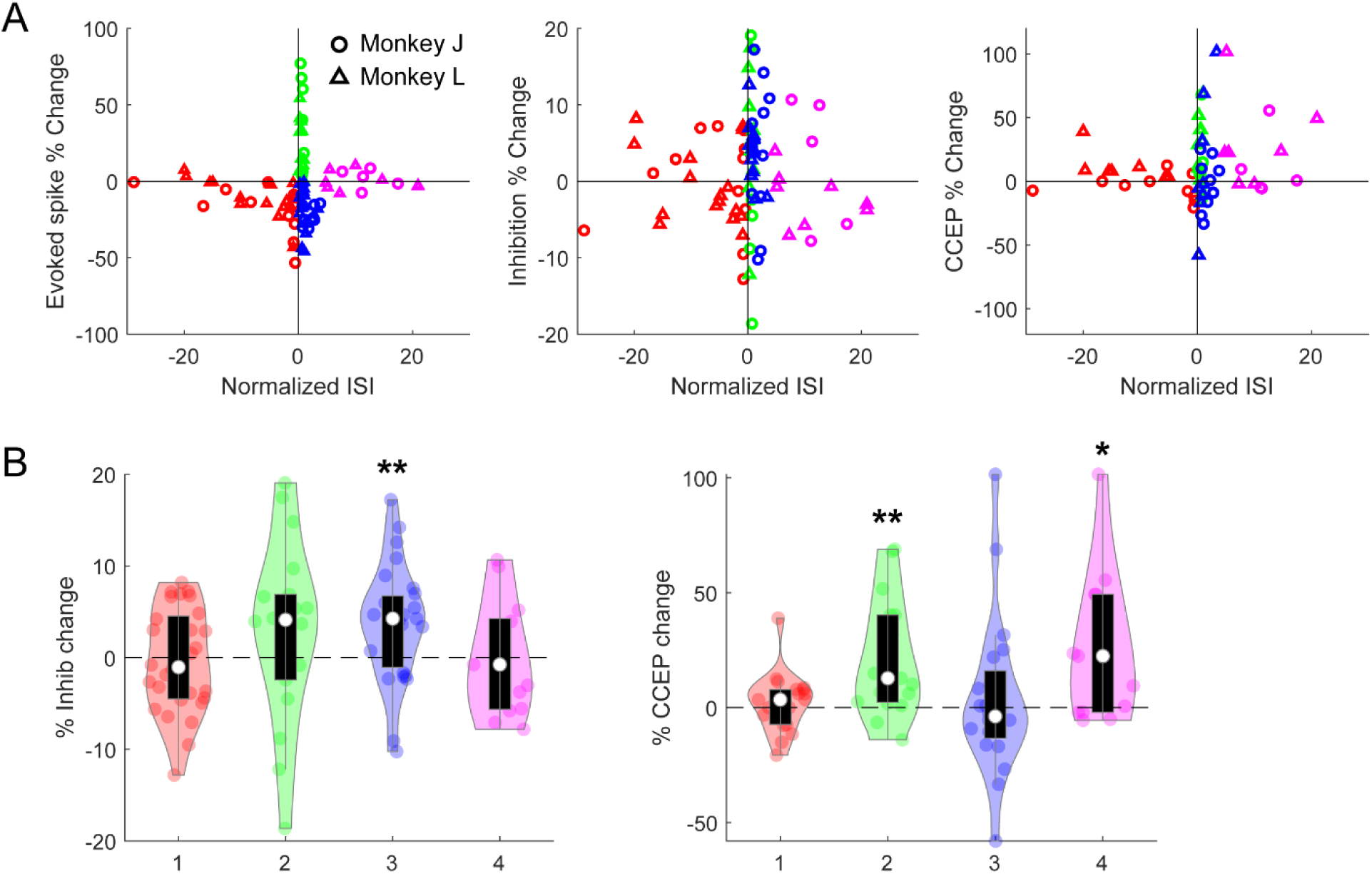
Comparisons in different ranges of ISIs. **A**. Four different regions of ISI: negative ISI, positive ES changes with ISI from 0 to 4, negative ES changes with ISI from 0 to 4, and ISI>4. Scatter plot of evoked spike probability change, inhibition duration change, and CCEP magnitude change shown with the 4 regions. **B**. Distributions of inhibition duration (left) and CCEP magnitude (right) changes in each region of ISI. (∗: p<0.05, ∗∗: p<0.01, Wilcoxon sign rank test) (0.004, 0.003, 0.03 in order from left to right)

Nevertheless, the measures may have a nonlinear relationship; the same magnitude of change in inhibition duration in one channel pair may manifest differently in another channel pair. Therefore, we simplified the comparisons by grouping the changes in each region rather than looking for a direct correlation. Figure 9C shows that the inhibition duration typically increased when evoked spike probability decreased, but only with positive ISIs. CCEPs, on the other hand, were likely to increase with positive ISIs, but not in sessions with decreased evoked spike probability (Figure 9C). These results suggest that the changes induced by conditioning are reflected in each measure and that the underlying mechanisms may be affecting one another.

### Persistence during the pre- and post-test epochs

We additionally determined whether the measures have baseline fluctuations over time during the pre-test epoch, and whether the changes we induced are consistent across the duration of the 10-minute post-test epoch. Figure 10 shows the aggregate z-scored changes in each measure over time, with significance from the first 10 seconds of the epoch denoted in red above. Evoked spike probabilities and inhibition duration were incredibly stable throughout the pre-test epoch, whereas the CCEP magnitude diminished slightly over time on average, though individual traces showed there are instances in which the magnitude increases over time. During the post-test epoch, the evoked spike probability had a slight but significant decrease over time and the inhibition duration had a slight but significant increase over time. The CCEP magnitude average seemed to be decreasing, but there was no statistical significance due to its variability.

**Figure 10.**
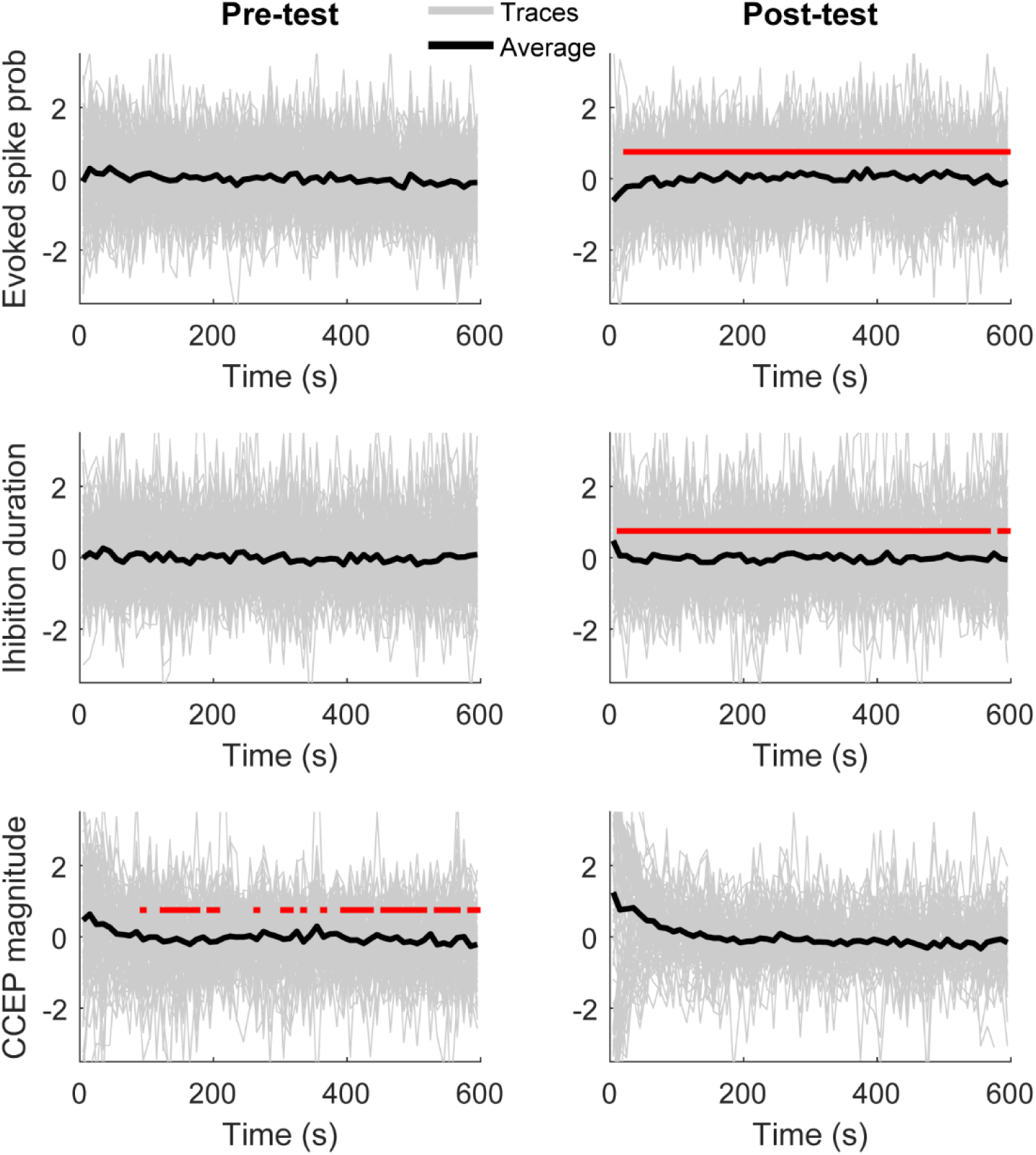
Measures over time. Average z-scored evoked spike probability (top), inhibition duration (middle), and CCEP magnitude (bottom) over time during the pre- and post-test epochs (left and right, respectively) using 10 second bins. Grey traces show individual sessions, black traces show the average, and the horizontal red bars show statistically significant difference from the first 10 seconds (p<0.05, Repeated measures ANOVA, Bonferroni correction post-hoc,).

However, as before, the changes that we observed during the post-test epoch may be specific to a range of ISIs. Using the four regions of ISIs previously defined, we further separated the changes in the measures over time (Figure 11). In sessions with negative ISIs (Figure 11, column 1) the inhibition duration change had a slight increase over time. This trend was reflected in the evoked spike probability as well but was not statistically significant. CCEP magnitude decreased over time on average but was highly variable and was not significant. In sessions with positive ISIs with increases in evoked spike probability (Figure 11, column 2) there was no consistent significant change over time in any measure. In sessions with positive ISIs with decreases in evoked spike probability (Figure 11, column 3) the evoked spike probably was initially low for the first 60 seconds that gradually increased to steady state over the course of the 10-minute epoch. This consistent trend was not reflected in any other measure. Finally, large positive ISIs (Figure 11, column 4) showed a decrease in CCEP magnitude over time.

**Figure 11.**
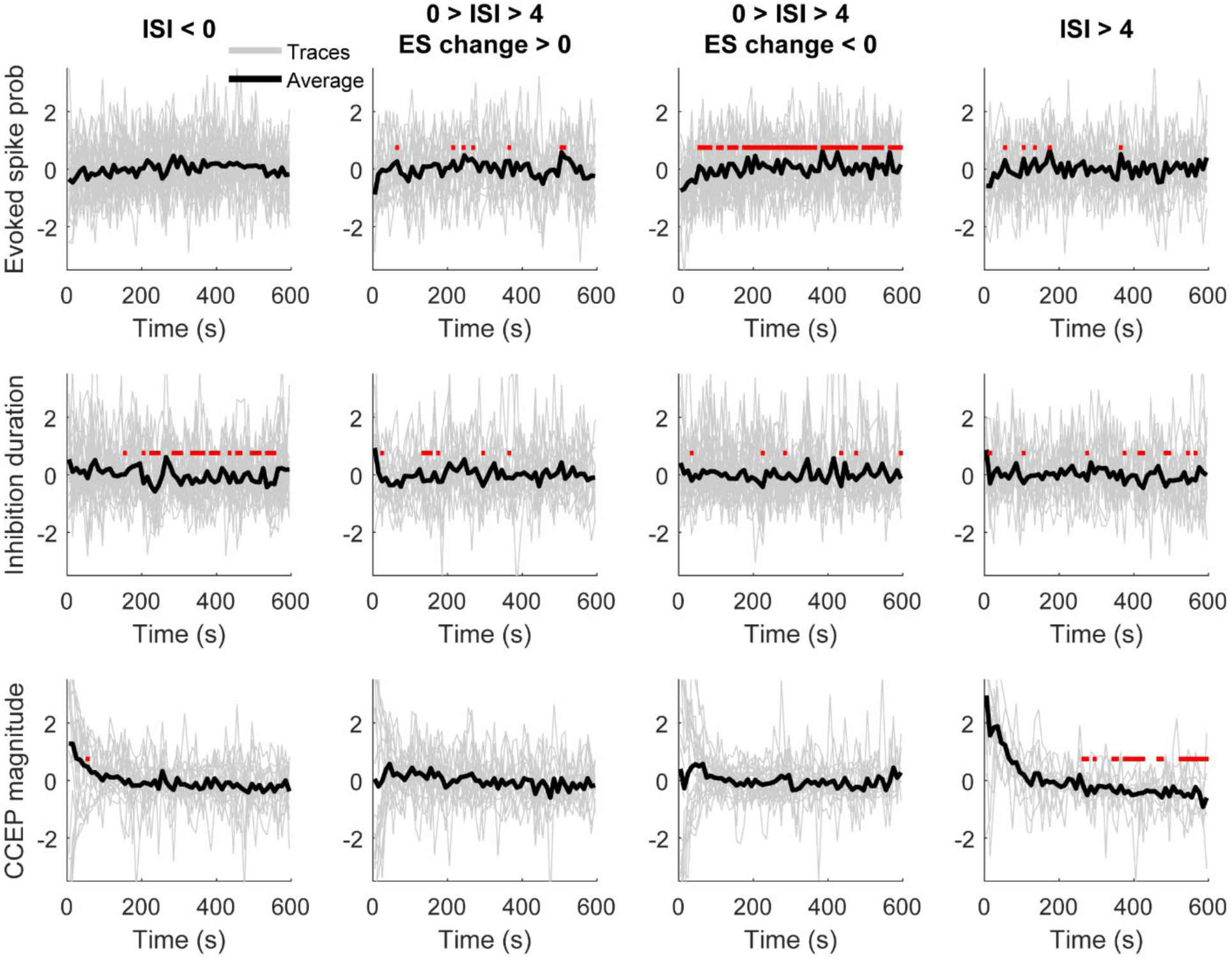
Measures over time during the post-test epoch by different regions of ISIs. Average z-scored evoked spike probability (top), inhibition duration (middle), and CCEP magnitude (bottom) over time during the post-test epochs for four different groups of normalized ISIs. Grey traces show individual sessions, black traces show the average, and the horizontal red bars show statistically significant difference from the first 10 seconds (p<0.05, Repeated measures ANOVA, Bonferroni correction post-hoc).

To assess whether the changes in the measures we initially observed were simply due to the averaging of short-term effects, we plotted the previous scatter plots using the first minute of the post-test period and again with the last minute of the post-test period (Figure 12). Changes in evoked spike probability were similar to the composite results when using the first minute, but the depression in positive ISIs was exaggerated. The depression was still present when only considering the last minute but was slightly diminished. The increase in probability of evoked spikes at positive ISIs showed a similar trend, being stronger during the first minute compared to the last minute. Interestingly, large positive ISIs resulted depression during the first minute that switched into being generally potentiation during the last minute.

**Figure 12.**
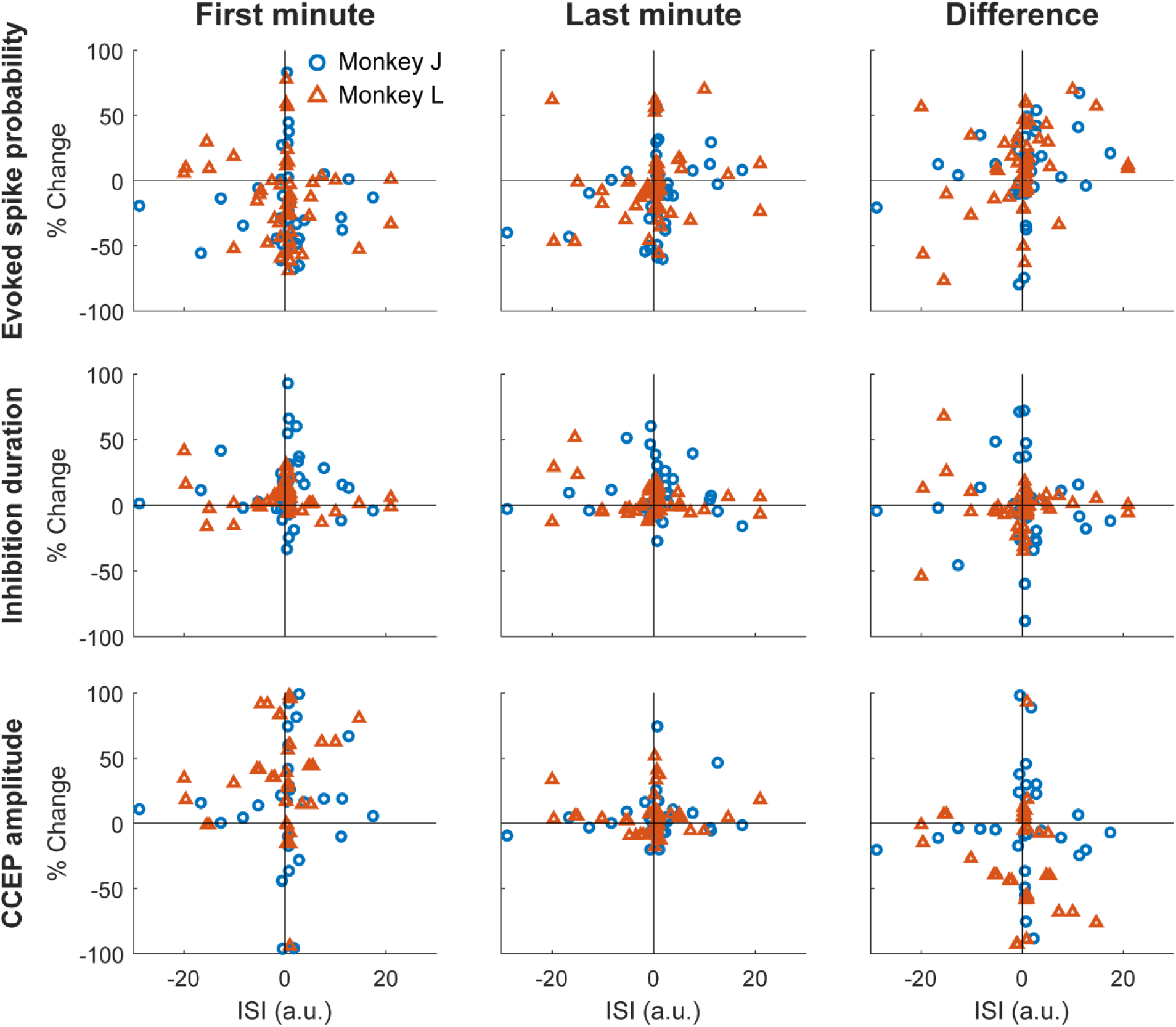
Changes in each measure during the first and last minute of the post-test epoch. Changes in evoked spike probability (top), inhibition duration (middle) and CCEP amplitude (bottom) with respect to normalized ISIs during the first (left) and last (middle) minute of the post-test epoch. The right column shows the difference between the last and first minute.

Changes in inhibition duration were more variable during the first minute. The only noticeable consistent change was the decrease during negative ISIs, which were mostly be contained within smaller ISIs. Changes in CCEPs generally had high magnitudes during the first minute that decreased by the last minute. This may be related to the overall decrease we see in CCEP magnitude over time even during the pre-test epoch before any conditioning.

### Control experiments

We performed experiments with six different control conditions: 1) Long delay – ISI of greater than ±100 ms, 2) Random stimulation – conditioning stimuli delivered randomly rather than time locked between the Pre and Post sites, 3) Subthreshold stimulation – 5 µA during conditioning shown to not evoked spikes, 4) No conditioning – no stimulation during conditioning, 5) Pre stimulation only – stimuli delivered only to the Pre site during conditioning, and 6) Post stimulation only – stimuli delivered only to the Post site during conditioning (Figure 13).

**Figure 13.**
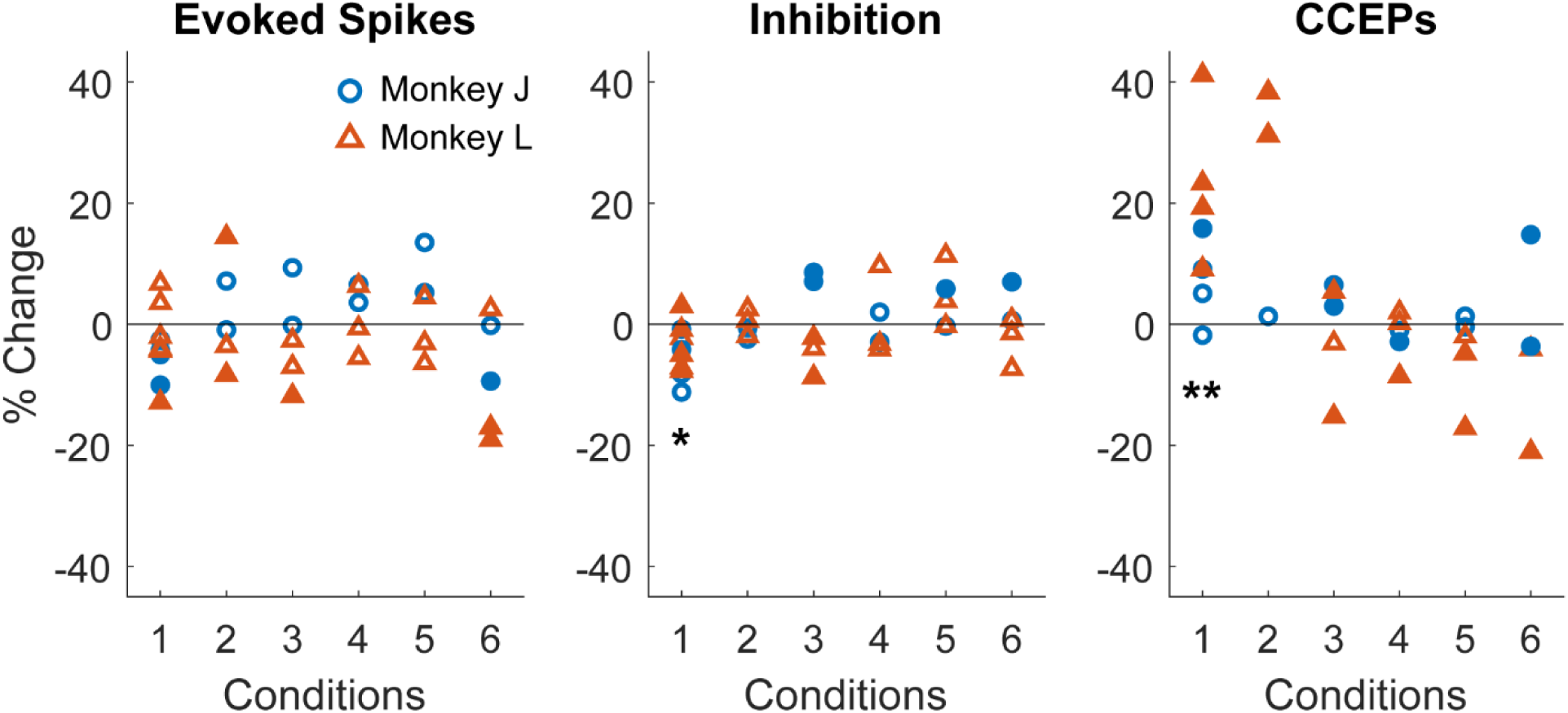
Controls. Percent change in evoked spike probability, inhibition duration, and CCEP magnitudes during different control experiment conditions: 1. Long delay, 2. Random stimulation, 3. Subthreshold stimulation, 4. No conditioning, 5. Pre-stimulation only, 6. Post-stimulation only. (∗: p<0.05, ∗∗: p<0.01, Wilcoxon sign rank test) (p=0.014 and 0.008).

Changes in evoked spike probability and inhibition duration were all below 20% under any control condition. Evoked spike probability did not have a consistent change under any control condition, but inhibition duration typically decreased during Long delay experiments. CCEPs showed much larger changes of up to over 40%, and typically increased during Long delay experiments.

We also analyzed whether there were changes in any measure during the post-test epoch for each control condition (Supplementary Figure 3). Although there was no statistically significant change over time in any condition, CCEP magnitudes typically decreased over time during Long delay and Random stimulation experiments.

## Discussion

### Summary of findings

Paired electrical stimulation in the primate motor cortex induced changes in connectivity between two sites. The probability of evoking spikes in the postsynaptic site when stimulating the presynaptic site deviated from classic STDP; positive ISIs less than 20 ms often resulted in depression. We observed that potentiation only occurred when the stimulus delivered to the Post site arrived before the timing of the evoked spike. The duration of the inhibitory response following the stimulus evoked spikes also did not reflect classic STDP, but typically increased when evoked spike probability decreased during positive ISIs. CCEPs also did not reflect classic STDP, resulting in some depression in positive ISI experiments, and typically increased when the evoked spike probability increased as well as with large positive ISIs. Direct comparisons between all three measures did not show any significant correlations.

Analyses of the measures over time showed that both evoked spike probability and inhibition duration were consistent during the pre-test epoch, but CCEPs decreased over time. Further analysis of the post-test epoch showed that evoked spike probabilities increased over time in positive ISIs that induced depression, inhibition duration increased over time in negative ISIs, and CCEPs decreased over time in large positive ISIs. Most control conditions showed that there were no large consistent changes in all measures, but inhibition duration decreased and CCEP magnitude increased during experiments with a long delay between the paired stimuli.

### Paired stimulation *in vivo* and single unit responses

Although delivering electrical stimulation to a cortical site activates excitatory fibers projecting to a neighboring site, it also activates fibers projecting to interneurons in the local circuitry that subsequently inhibit the recorded principal cells (Butovas et al., 2006; Logothetis et al., 2010; Yun et al., 2022). Thus, improper timing of stimulation delivered to the Post site may enhance the inhibitory activity rather than the firing of the spike (Figure 14).

**Figure 14.**
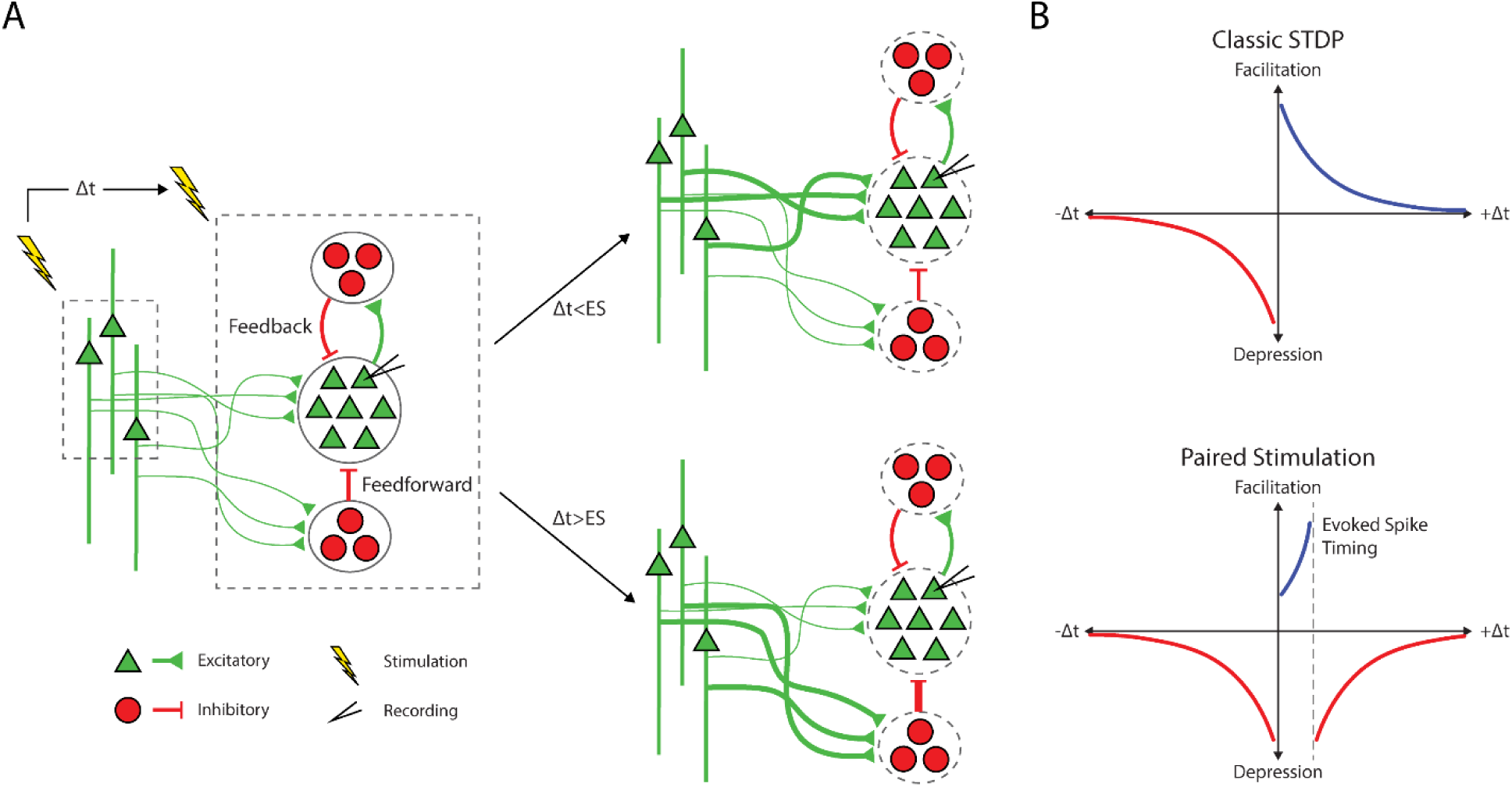
Mechanisms. **A**. Stimulation at the Pre site activates excitatory horizontal fibers, activating the recorded principal neurons and the local feedforward and feedback inhibitory circuitry. Paired stimulation strengthens the projections to the principal cells if the stimulation to the Post site arrives before the timing of the evoked spike, whereas it strengthens the feedforward inhibitory pathway if the stimulation arrives after the evoked spike. **B**. Schematic of classic STDP (top) compared to what we observed with paired stimulation and single neuron response (bottom).

Our findings support this hypothesis as facilitation is only present when the stimulation of the Post site occurs before the timing of the evoked spike. In addition, the inhibition duration typically lengthens in positive ISI experiments that resulted in fewer stimulus evoked spikes, which further suggests the inhibitory circuitry becomes stronger or more active. The variability in the changes is both likely due to the initial evoked spike probability as well as the fact that inhibitory activity may be occurring as soon as the excitatory response due to their high connectivity and large number of horizontal projections in layer 2/3 (Adesnik & Scanziani, 2010; Isaacson & Scanziani, 2011).

The lack of symmetry in the distribution of changes in evoked spike probabilities is likely due to the specificity of the measure – we are measuring single unit responses in the Post-site and stimulating the Pre-site – resulting in non-specific changes when using negative ISIs. Following STDP, negative ISIs would result in excitatory projections from the Pre-site to the Post-site being weakened and projections from the Post-site to the Pre-site being strengthened. However, there is variability in the magnitude of changes in specific synapses and no guarantee of uniform changes on the synapses to and from the recorded stimulus evoked spike, which may explain the variability present in the strength of depression in negative ISI conditions.

Our findings do not contradict STDP as the changes are likely STDP mediated. The magnitude and direction of change was heavily dependent on the ISI and lasted for at least 10 minutes. Instead, the results suggest that stimulus induced plasticity in vivo is much more complex than previously thought, and the interactions within and between local circuitry must be taken into account.

Additionally, analysis of evoked spike probability and inhibition duration over time showed that the measures fluctuated over the 10-minute post-test epoch, especially in experiments that saw a decrease in the probability with evoked spikes with positive ISIs. As the curves seem to return to equilibrium, one theory could be homeostatic plasticity, but synaptic scaling typically occurs over the timescale of hours or days (G. Turrigiano, 2012; G. G. Turrigiano, 2008). Another explanation could be that there are lingering effects of neurotransmitters receptors which can take time to return to baseline. However, the time constants of post-synaptic current of both glutamate and GABA receptors are within 100’s of milliseconds, whereas the changes over time we observed lasted for 10’s of seconds (Bettler et al., 2004; Thomson & Destexhe, 1999). The timescale does match short-term plasticity (Zucker & Regehr, 2002); however, short-term plasticity is typically induced by open loop stimulation, and as such we should observe the same changes over time regardless of the ISI as well as in control experiments. Thus, we may be observing some other form of regulating plasticity.

### Comparison with CCEPs

CCEPs had similar changes as evoked spike probability, but with much higher variability. Although CCEPs are directly indicative of neural responses, they are a measure of population activity which may detract from the changes induced by the conditioning (Keller et al., 2014; Prime et al., 2020; Vincent et al., 2017). This is confirmed in our study in which CCEPs are independent of the stimulus evoked spikes, as they are a result of greater spatial integration. We additionally observed more variable changes in CCEPs due to conditioning relative to the ISI, similar to a previous experiment studying paired stimulations with CCEPs in which a limited number of channel pairs could be induced with STDP (Seeman et al., 2017).

Intracortical CCEPs, especially when used as a measure of connectivity, are most commonly measured using the peak-to-peak amplitude under the assumption that it reflects synchronized firing of spikes – more spikes firing as a result of stimulation result in a larger amplitude CCEP (Seeman et al., 2017; Zanos et al., 2018). However, this interpretation of CCEPs suggests that proper timing of stimulation to arrive at or before the trough should reliably induce potentiation, which was not observed in our study. Instead, CCEPs may be reflective of excitatory responses in conjunction with inhibitory responses and possibly involve complex spatiotemporal summation (Keller et al., 2014; Vincent et al., 2017). Different measures suggested from previous literature, such as root-mean squared and area under the curve, did not provide more consistent results (Prime et al., 2020).

Additionally, we observed that CCEPs typically decrease over time during the pre-test epoch before any conditioning has been applied, also observed in previous studies when using low frequency (<20 Hz) stimulation (Goldring et al., 1961; Vincent et al., 2016, 2017). This may explain the findings in a previous study using CCEPs to assess the effects of paired stimulation which showed a global increase in CCEPs following conditioning (Seeman et al., 2017). The decrease in CCEP amplitude has been attributed to distortions from a saturated amplifier, which is not applicable in our study as the shape of the CCEP does not change over time, or to a desynchronization of cortical networks (Vincent et al., 2016). However, during the post-test epoch we also observed large increases over time, and control experiments also did not provide consistent CCEPs. Additional studies are necessary to fully understand why CCEPs change drastically over time.

Although CCEPs may be a useful metric for assessing the presence of connectivity, they may not be a simple metric for measuring the strength of and changes in connectivity. There is a need to better understand the neural mechanisms of invoking CCEPs, and the implications behind the timings and amplitudes of troughs and peaks, especially with how variable CCEPs are between different stimulus intensities and depth of recording.

### Implications to *in vivo* plasticity paradigms

Plasticity paradigms *in vivo* often require high amplitudes or continuous conditioning for long periods of time for changes in connectivity (Jackson et al., 2006; Rebesco & Miller, 2011; Seeman et al., 2017; Yazdan-Shahmorad et al., 2018). However, our experiment showed consistent changes with just 10 minutes of low amplitude paired stimulation. This is likely attributed to two reasons: the measure we were using is very specific and allowed us to assess smaller changes, and the timing of our stimulation was very exact relative to the stimulus response. Assessing the connectivity with finer details and tailoring stimulation paradigms to the responses may allow for more effective stimulation paradigms that can be reasonable in a clinical setting.

However, even within the cortex there could be discrepancies in stimulus responses and induced plasticity in different cortical regions or even cortical layers. Our study was in the motor cortex, which is notorious for its high interconnectivity, and in layer 2/3 which has more inhibitory horizontal projections. Studies exploring changes in connectivity in different layers, especially layer 5 with its large pyramidal cells, or different cortical regions, like the visual system and its unidirectional information flow, can confirm whether our findings are generalizable.

## Conclusions

We applied paired stimulation to the motor cortex of awake primates using a finer temporal gradient of ISIs than tested before. 10 minutes of conditioning generated consistent changes in the responses of single units dependent on the ISI. Contrary to classic STDP, very short, positive ISIs resulted in depression, likely due to potentiation in the local inhibitory circuitry, suggesting that the dynamics of plasticity in vivo may have a much finer timescale than previously thought due to the interconnected nature of excitatory and inhibitory circuitry. CCEPs showed similar but much more variable results compared to single unit responses, indicating that they may not be the ideal measure for changes in connectivity strength. Better understanding underlying dynamics of stimulus responses and connectivity metrics could lead to more effective plasticity paradigms.

## Acknowledgments

We thank Larry Shupe for programming and software support and Rebekah Schaefer and Becky Burch for assistance with animal care, handling, training, and surgery. We also thank Irene Rembado and Rajesh Rao for helpful discussion. This work was supported by the National Institutes of Health (NS012542, RR00166, and NS118781) and the National Science Foundation (EEC-1028725).

## Supplementary Figures

**Supplementary Figure 1.**
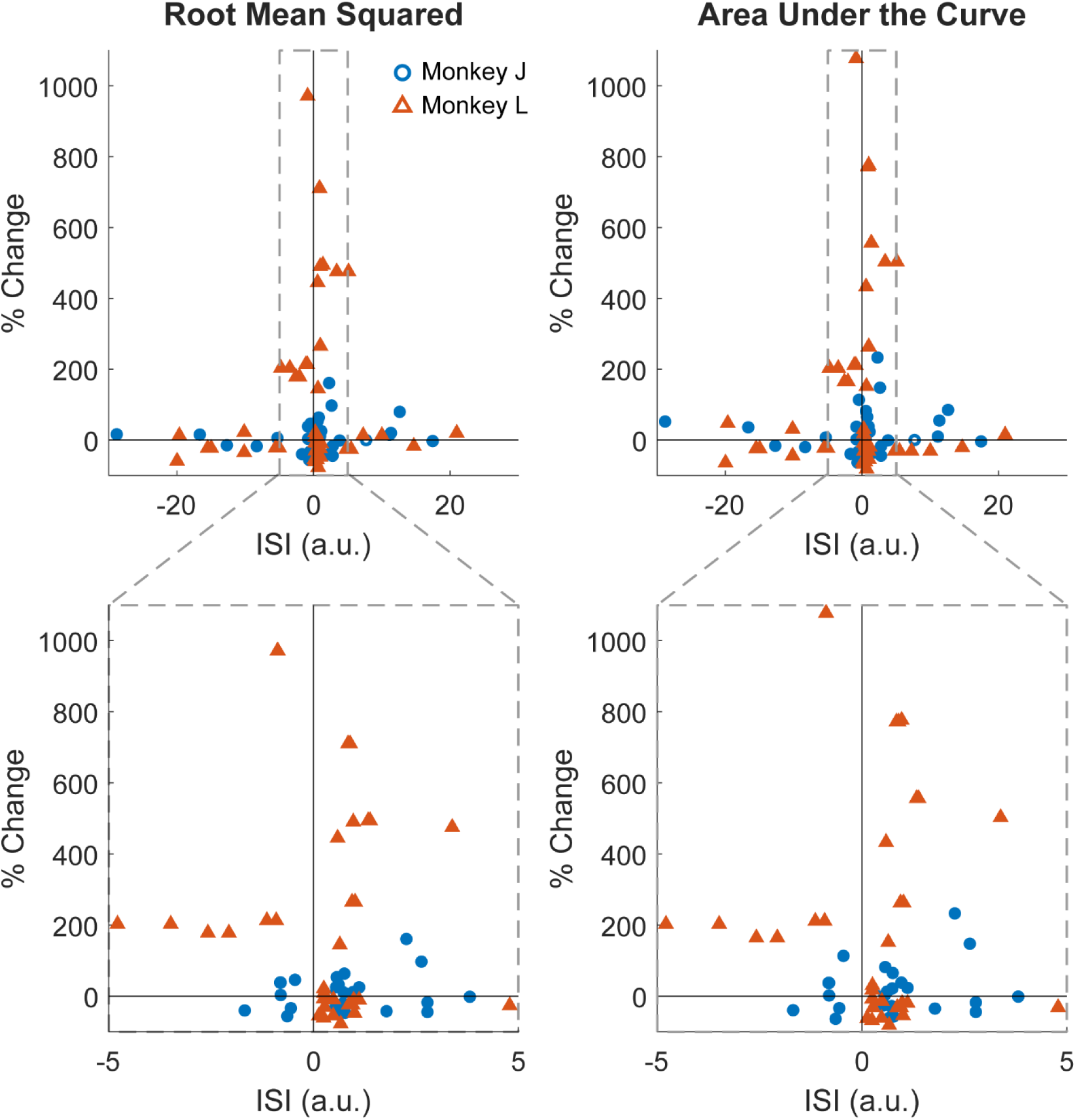
Different CCEP measures. Changes in CCEPs before and after conditioning calculated using root-mean squared (left) or area under the curve (right) with respect to normalized ISIs.

**Supplementary Figure 2.**
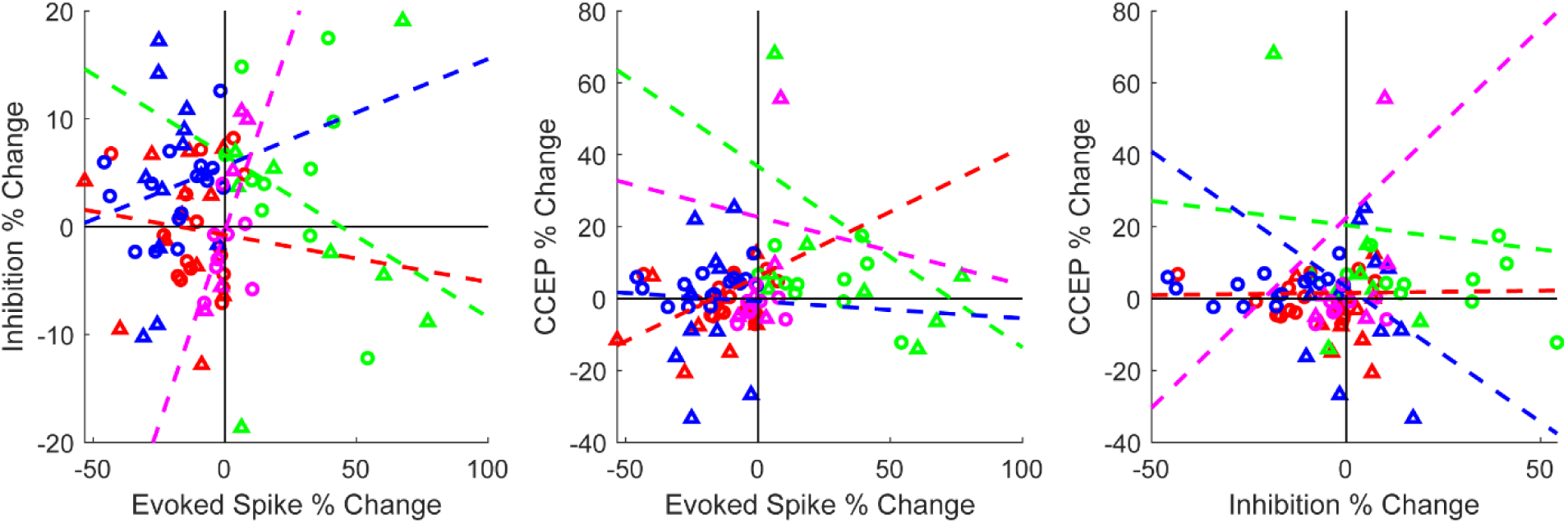
Correlations for each region of ISI. Scatter plots and best fit lines of each ISI region for every pair of measures. There is no statistically significant correlation between any measure in any region.

**Supplementary Figure 3.**
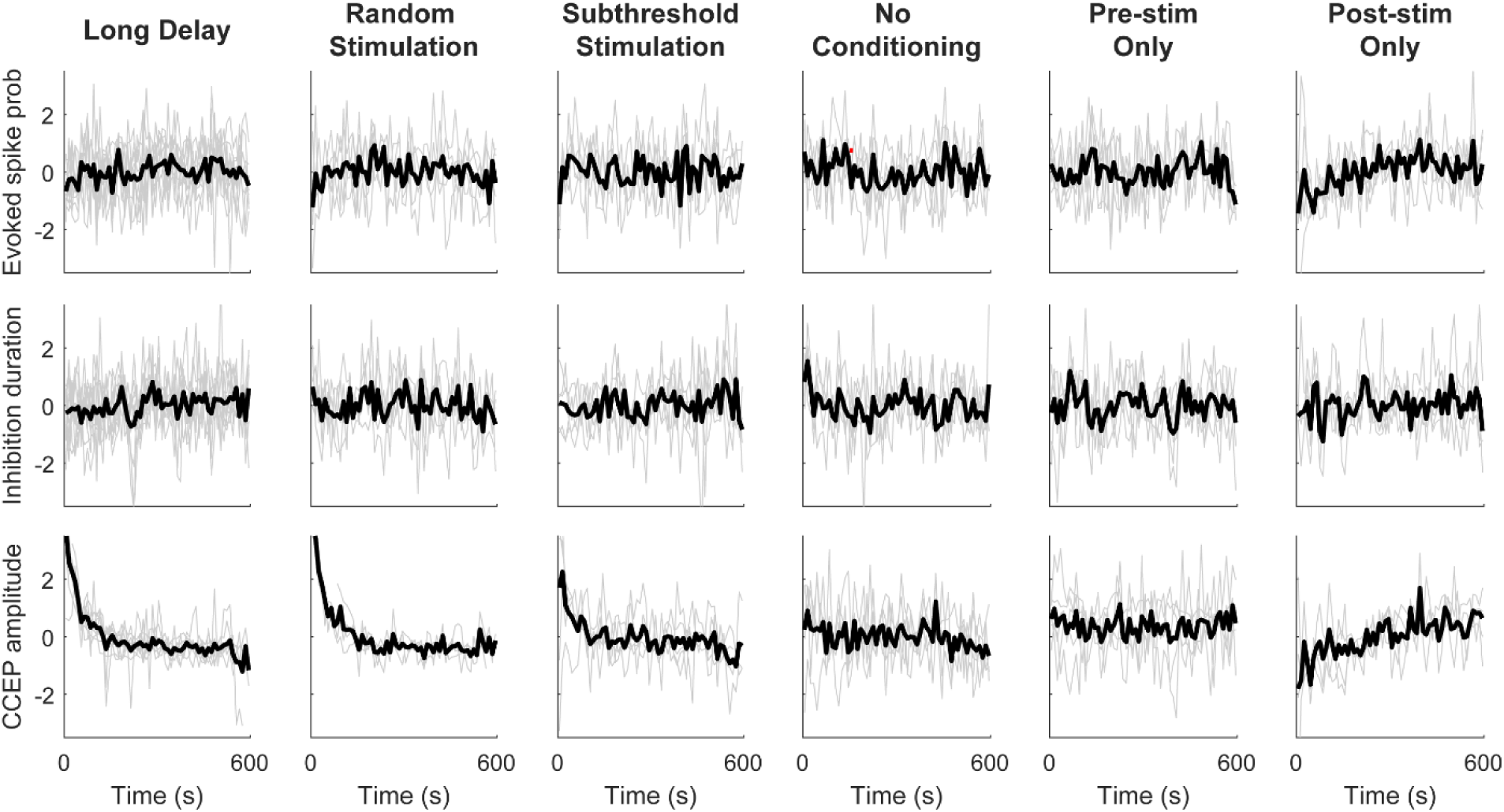
Controls over time. Individual session traces (grey) and averages (black) of z-scored evoked spike probability (top), inhibition duration (middle), and CCEP amplitude (bottom) during the 10-minute post-test epoch for each control condition. There is no statistical significance between the first 10 seconds to the rest of the epoch in any measure (repeated measures ANOVA, p<0.05).

